# Optimal Recovery of Missing Values for Non-negative Matrix Factorization

**DOI:** 10.1101/647560

**Authors:** Rebecca Chen, Lav R. Varshney

## Abstract

We extend the approximation-theoretic technique of optimal recovery to the setting of imputing missing values in clustered data, specifically for non-negative matrix factorization (NMF), and develop an implementable algorithm. Under certain geometric conditions, we prove tight upper bounds on NMF relative error, which is the first bound of this type for missing values. We also give probabilistic bounds for the same geometric assumptions. Experiments on image data and biological data show that this theoretically-grounded technique performs as well as or better than other imputation techniques that account for local structure.

## I. Introduction

MATRIX factorization is commonly used for clustering and dimensionality reduction in computational biology, imaging, and other fields. Non-negative matrix factorization (NMF) is particularly favored by engineers and biologists because non-negativity constraints preclude negative values that are difficult to interpret in biological processes [2], [3]. A recent tutorial article highlighted the interpretability and identifiability (or model uniqueness) of NMF, both of which are valuable for practical applications [4]. Without constraints on the factor matrices, latent factors are non-unique, but requiring non-negativity guarantees model uniqueness under certain assumptions. Furthermore, experimental results demonstrate that the latent factors are intuitive given the data [4]. In signal processing and statistical learning, NMF has been used for speech and audio separation, medical imaging, community detection, and topic modeling. In biology, NMF of gene count matrices can discover cell groups and lower-dimensional manifolds (latent factors) describing gene count ratios for different cell types. Due to channel noise, incomplete survey data, or biological limitations, however, data matrices are usually incomplete and matrix imputation is often necessary before further analysis [3]. In particular, Stein-O’Brien et al. argue that “newer MF algorithms that model missing data are essential for [single-cell RNA sequence] data” [5].

Imputation accuracy is commonly measured using root mean-squared error (RMSE) or similar error metrics. However, Tuikkala et al. argue that “the success of preprocessing methods should ideally be evaluated also in other terms, for example, based on clustering results and their biological interpretation, that are of more practical importance for the biologist” [6]. Here, we specifically consider imputation performance in the context of NMF and describe NMF error rather than RMSE (or some other prediction error) of imputed matrices. Previous analyses of information processing algorithms with missing data have considered high-dimensional regression [7] and subspace clustering [8]. Missing values for NMF have also been studied for the application of stock price prediction, but previous approaches lack theoretical guarantees [9].

Data often exhibits local structure, e.g., different groups of cells follow different gene expression patterns. Information about local structure can be used to improve imputation. We introduce a new imputation method based on *optimal recovery*, an approximation-theoretic approach for estimating linear functionals of a signal [10]–[12] previously applied in signal and image interpolation [13]–[15], to perform matrix imputation of clustered data. Characterizing optimal recovery for missing value imputation requires a new geometric analysis technique. Previous work on missing values take a statistical approach rather than a geometric one.

Our contributions include:

- A computationally efficient imputation algorithm that performs as well as or better than other modern imputation methods, as demonstrated on hyperspectral remote sensing data and biological data;
- A tight upper bound on the relative error of downstream analysis by NMF. This is the first such error bound for settings with missing values; and
- A probabilistic bound on NMF error after imputation.

The remainder of the paper is organized as follows. In Section II, we give background on missing data mechanisms, imputation algorithms, and NMF. In Section III, we introduce optimal recovery and apply it to NMF. In Section IV, we present an algorithm for optimal recovery imputation of clustered data and give a deterministic upper bound on algorithm performance. In Section V we give a probabilistic bound on the performance of our algorithm. In Section VI we give experimental results for both synthetic and real data, and we conclude in Section VII.

## II. Background

In this section, we describe the relationships between missingness patterns and the underlying data, which are referred to as *missing data mechanisms*. We then discuss prior work on practical imputation algorithms, and we present NMF as an analysis technique that is commonly performed after imputation.

### A. Missingness mechanisms

Rubin originally described three mechanisms that may account for missing values in data: missing completely at random (MCAR), missing at random (MAR), and missing not at random (MNAR) [16]. When data is MCAR, the missing data is a random subset of all data, and the missing and observed values have similar distributions [17]. The MCAR condition is described in (1), where X refers to observed variables and Y refers to missing variables. This may occur if a researcher forgets to collect certain information for certain subjects, or if certain data samples are collected only for a random subset of test subjects. When data is MAR, the distribution of missing data is dependent on the observed data, (2). For example, in medical records, patients with normal blood pressure levels are more likely to have missing values for glucose levels than patients with high blood pressure. When data is MNAR, the distribution of missing data is dependent on the unobserved (missing) data, (3). For example, people with very high incomes may be less likely to report their incomes.

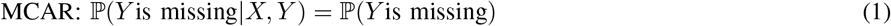

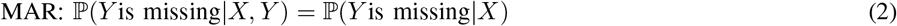

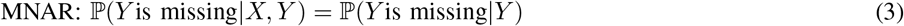

It is important to understand the missingness mechanism when analyzing data. When data is MCAR, the statistics of the complete cases (data points with no missing observations) will represent the statistics of the entire dataset, but the sample size will be much smaller. If data is MAR or MNAR, the complete cases may be a biased representation of the dataset. One can also estimate statistics such as means, variances, and covariances based on all available non-missing observations of a variable [18]. Then the sample size reduction may be less severe for certain variables. However, this may be a biased representation of the entire dataset when data is MAR or MNAR, and there is the additional problem of inconsistent sample sizes. Although some research has been done on MNAR imputation, this is generally a difficult problem, and most imputation methods assume the MAR or MCAR model.

Ding and Simonoff argue that missingness mechanisms are more nuanced than the three basic categories described by Rubin [19]. They claim that missingness can be dependent on any combination of the missing values, the observed predictors, and the response variable (e.g. a category label). In cases where the missingness pattern contains information about the response variable, the missingness is *informative* [20]. Ghorbani and Zou leverage informative missingness, using the missingness patterns themselves as an additional feature for data classification [21].

### B. Imputation algorithms

Imputation is often necessary before specific downstream analysis, such as clustering or manifold-finding for classification. Two main categories of imputation are single imputation, in which missing values are imputed once, and multiple imputation, in which missing values are imputed multiple times with some built-in randomness. The variance in the multiple imputations of each missing observation reflects the uncertainty of the estimates, and *all* imputed datasets are used in the downstream analysis, which increases statistical power.

#### 1) Single imputation

One of the simplest imputation techniques is *mean imputation*. Missing values of each variable are imputed with the mean value of that variable. Since all missing observations of a variable are imputed with the same value, variance is reduced, and other statistics may be skewed in the MAR and MNAR cases. The reduced variability in the imputed variable also decreases correlation with other variables [22].

In *regression imputation*, a variable of interest is regressed on the other variables using the complete cases. Imputation puts points with missing values directly on the regression line. This method also underestimates variances, but it overestimates correlations. *Stochastic regression* attempts to add the variance back by distributing imputed points above and below the regression line using a normal distribution.

Bayesian imputation approaches also exist, including *Bayesian PCA* [23] and *maximum likelihood imputation* [24]. Bayesian methods are theoretically sound and assume that data samples are generated from some underlying joint distribution. In practice, these methods require numerical algorithms such as the Markov chain Monte Carlo (MCMC) method, which may be prohibitively time-consuming for large datasets.

#### 2) Multiple imputation

Multiple imputation attempts to preserve the variance/covariance matrix of the data. Several imputations are randomly generated, resulting in multiple complete datasets. Imputed datasets are then analyzed and results are pooled. The different imputations introduce variance into the data, but the variance may still be an underestimate since the imputations assume correlation between the variables. One popular algorithm for multiple imputation is *multiple imputation by chained equations* (MICE) [25]. While MICE does not have the theoretical backing that maximum likelihood imputation has, MICE is flexible and can accommodate known interactions and independencies of real-world datasets [26]. A stepwise regression can be performed so that the missing variable is regressed on the best predictors.

### C. Imputation with clustered data

When the underlying data is clustered, a data point should be imputed based on its cluster membership. Local imputation approaches outperform global ones when there is local structure in data. Global approaches generally perform some form of regression or mean matching across all samples [25], [27], whereas local approaches group subsets of similar samples. Popular imputation algorithms that utilize local structure include k-nearest neighbors (kNN), local least squares (LLSimpute), and bicluster Bayesian component analysis (biBPCA) [28]–[30]. The kNN imputation method finds the k closest neighbors of a sample with missing values (measured by some distance function) and fills in the missing values using an average of its neighbors. LLSimpute uses a multiple regression model to impute the missing values from k nearest neighbors. Rather than regressing on *all* variables, biBPCA performs linear regression using biclusters of a lower-dimensional space, i.e. coherent clusters consisting of correlated variables under correlated experimental conditions. Delalleau et al. develop an algorithm to train Gaussian mixtures with missing data using expectation-maximization (EM) [31]. By itself, MICE does not address clusters, but cluster-specific (group-wise) regression can be performed [32].

Tuikkala et al.’s clustering results on cDNA microarray datasets showed that “even when there are marked differences in the measurement-level imputation accuracies across the datasets, these differences become negligible when the methods are evaluated in terms of how well they can reproduce the original gene clusters or their biological interpretations” [6]. They use the Average Distance Between Partition (ADBP) to calculate clustering error, and they show that BPCA, LLS, and kNN give similar clustering results. Chiu et al. find that LLS-like algorithms performed better than kNN-like algorithms in terms of downstream clustering accuracy (measured using cluster pair proportions) [33]. De Souto et al. evaluate whether the effect of different imputation methods on clustering and classification are statistically significant [34]. They remove all genes with more than 10% missing values and compare classification using the corrected Rand index. They find that simple methods such as mean and median imputation perform as well as weighted kNN and BPCA.

### D. Non-negative matrix factorization (NMF)

After imputation, downstream analysis such as NMF can be performed on data. Donoho and Stodden interpret NMF as the problem of finding cones in the positive orthant which contain clouds of data points [35]. Liu and Tan show that a rank-one NMF gives a good description of near-separable data and provide an upper bound on the relative reconstruction error [36]. Given that gene and protein expression data is often linearly separable on some manifold- or high-dimensional space [37], the bound given by rank-one NMF is valid. We extend these ideas to data with missing values and, for the first time, bound performance of downstream analysis of imputation. Loh and Wainwright have previously bounded linear regression error of data with missing values [7], but they do not consider imputation, and their proof is based on modifying the covariance matrix when data is missing. Tan and Févotte consider NMF with missing values, but they replace missing with zeros instead of performing imputation, and they do not provide error bounds [9]. In addition, their NMF algorithm requires hyperparameter tuning and makes some probabilistic assumptions on the data. Our proof is based on the geometry of NMF and is parameter-free. The nonnegativity of NMF lends itself to a geometric interpretation, which we describe in Section III.

## III. Optimal recovery

In this section, we introduce our new approach to imputation based on approximation-theoretic ideas. Suppose we are given an unknown signal *v* that lies in some signal class *C_k_*. The optimal recovery estimate 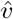 minimizes the maximum error between **V** and all signals in the feasible signal class. Given well-clustered non-negative data **V**, we impute missing samples in V so the maximum error is minimized over feasible clusters, regardless of the missingness pattern.

### A. Application to clustered data

Let 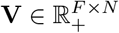 be a matrix of *N* sample points with *F* observations (*N* points in *F*-dimensional Euclidean space). Suppose the *N* data points lie in *K* disjoint clusters *C_k_* (where *k* = 1, 2,…, *K*), and that these clusters are compact, convex spaces (e.g., the convex hull of the points belonging to *C_k_*).

Now suppose there are missing values in **V**. Let Ω ∈ {0, 1}*^F×N^* be a matrix of indicators with Ω*_ij_* = 1 if *v_ij_* is observed and 0 otherwise. We make no assumptions on the missingness pattern, such as MCAR or MAR because we take a geometric approach rather than a statistical one. We define the projection operator of a matrix **Y** onto an index set Ω by

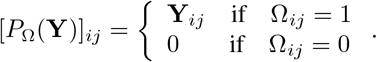

We use the subscripted vector (·)*_fo_* to denote fully observed data points (columns), or data points with no missing values, and we use the subscripted vector (·)*_po_* to denote partially observed data points. We use a subscripted matrix (·)*_fo_* or (·)*_po_* to denote the set of all fully observed or partially observed data columns in the matrix.

We can impute a partially observed vector *v_po_* by observing where its observed samples intersect with the clusters *C*_1_,…,*C_k_*. Let the *missing values plane* be the restriction set over 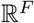 that satisfies the constraints on the observed values of *v_po_*. We call this intersection the *feasible set W*:

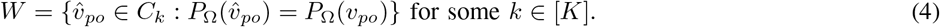

Fig. 1 illustrates the feasible set of a three-dimensional vector with two missing samples when the signal class (convex space containing samples from *k*th cluster) covers an ellipsoid. If the signal had only one missing sample, the feasible set would be a line segment.

**Fig. 1.**
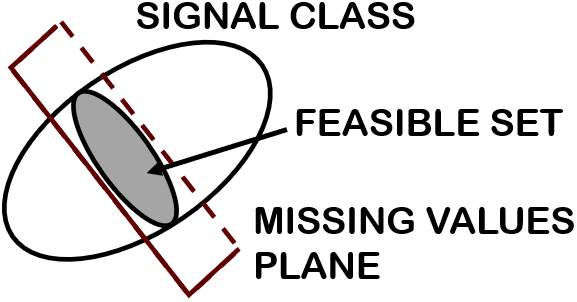
Feasible set of estimators.

All *k* for which (4) is satisfied are possible clusters from which the true *v* originated. Since *W* cannot be empty, there must be at least one *C_k_* that has non-empty intersection with the set of all points satisfying the *P*_Ω_(*v_po_*) constraint. The optimal recovery estimator 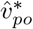 minimizes the maximum error over the feasible set of estimates:

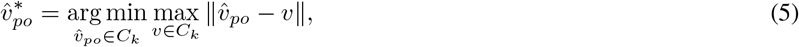

where || · || denotes some norm or error function. If we use the ∞-norm, 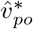 is the Chebyshev center of the feasible set. If *W* contains estimators belonging to more than one *C_k_*, *W* can be partitioned into *K* disjoint sets, *W_k_*, defined as

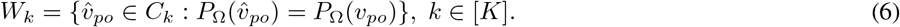

Feasible clusters are those for which *W_k_* is not empty, and we can find (5) over the *C_k_* for which the corresponding *W_k_* covers the largest volume: *k* = arg max*_k_* |*W_k_*|.

### B. Application to non-negative matrix factorization

Let 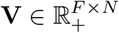 be a matrix of *N* sample points with *F* non-negative observations. Suppose the columns in **V** are generated from *K* clusters. There exist 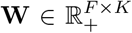 and 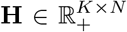 such that **V** = **WH**. This is the NMF of **V** [38]. We use the conical interpretation of NMF [35], [36], described as follows.

Suppose the *N* data points originate from *K* cones. We define a circular cone *C*(*u, α*) by a direction vector *u* and an angle *α*:

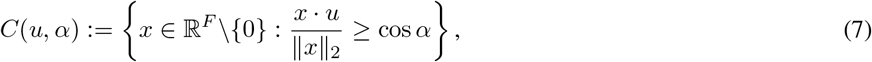

or equivalently,

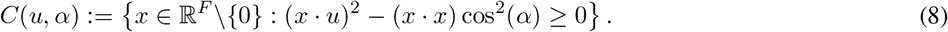

In a three-dimensional space, this conical hull is sometimes called an ice cream cone [4]. We truncate the circular cones to be in the non-negative orthant *P* so that we have *C*(*u, α*) ∩ *P*. We can consider *u_k_* to be the dictionary entry corresponding to *C_k_* and all *x*’s belonging to *C_k_* as noisy versions of *u_k_*. We call the angle between cones *β_ij_* := arccos(*u_i_* · *u_j_*). Assume the columns of **V** are in *K* well-separated cones, that is,

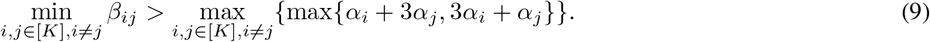

This implies that the distance between any two points originating from the same cluster is less than the distance between any two points in different clusters, which is a common assumption used to guarantee clustering performance [36], [39], [40] (see Fig. 2). We can then partition **V** into *k* sets, denoted **V***_k_* := {**v***_n_* ∈ *C_k_* ∩ *P*}, and rewrite **V***_k_* as the sum of a rank-one matrix **A***_k_* (parallel to *u_k_*) and a perturbation matrix **E***_k_* (orthogonal to *u_k_*). For any vector **z** ∈ **V***_k_*, **z** = ||**z**||_2_(cos *β*)*u_k_* + **y**, where ||**y**||_2_ = ||**z**||_2_(sin *β*) ≤ ||**z**||_2_(sin *α_k_*). We use this rank-one approximation to find error bounds [36] (see Fig. 3).

**Fig. 2.**
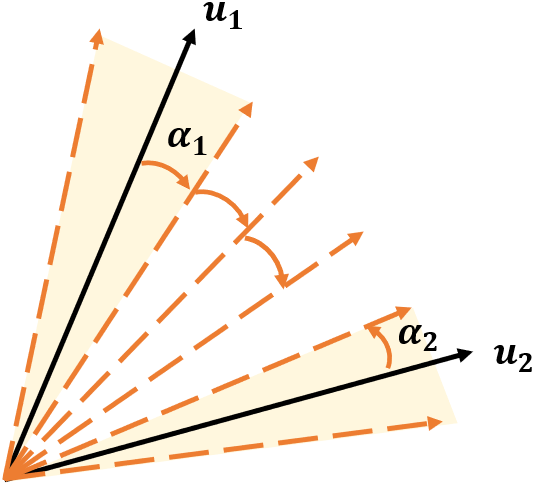
Geometric assumption for greedy clustering.

**Fig. 3.**
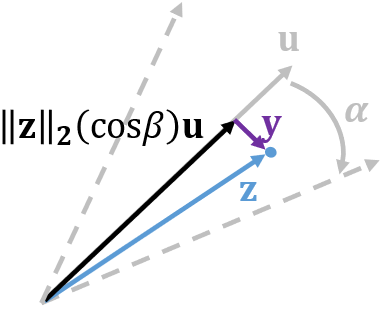
Decomposition of vectors in a circular cone.

If **V** contains missing values, we can use the optimal recovery estimator to impute **V**. Assuming the columns in **V** come from *K* circular cones defined as (7), there is a pair of factor matrices 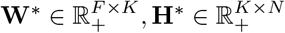, such that

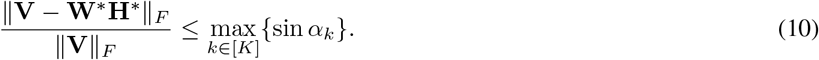

Since the error is bounded by sin *α_k_*, we choose our optimal recovery estimator to minimize *α_k_*. This is equivalent to maximizing the inequality in (8):

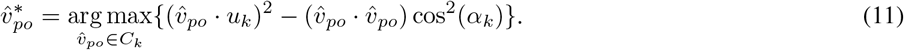

We can solve (11) analytically using the Lagrangian with known values of *v_po_* as equality constraints. We can also solve (11) numerically using projected gradient descent.

Generally, *u_k_* is not known beforehand, but we can find *u_k_* given *W_k_*. Given an ellipse in 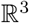, we reconstruct its cone by drawing lines from its limit points to the origin. Then it is straightforward to find the center of the cone. (Note that while this volume minimization problem is NP-hard, there are efficient and accurate algorithms when certain assumptions are met, which have been used with NMF [4].) Liu and Tan propose the following optimization problem (in the absence of missing values) over the optimal size angle and basis vector for each cluster [36]. We write the data points in each cluster as 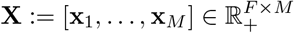 where 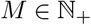:

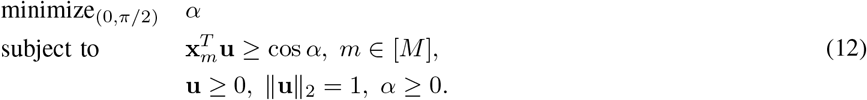

Of course, we also do not know *C_k_* or *W_k_*, so we use a clustering algorithm to find the vectors belonging to each *C_k_* (see Sec. IV).

## IV. Algorithm and error bound

Now we consider clustering and NMF with missing values. If the geometric assumption (9) holds, a greedy clustering algorithm [36, Alg. 1] returns the correct clustering of fully observed data. Here we show that a greedy algorithm also guarantees correct clustering of partially observed data under certain conditions.

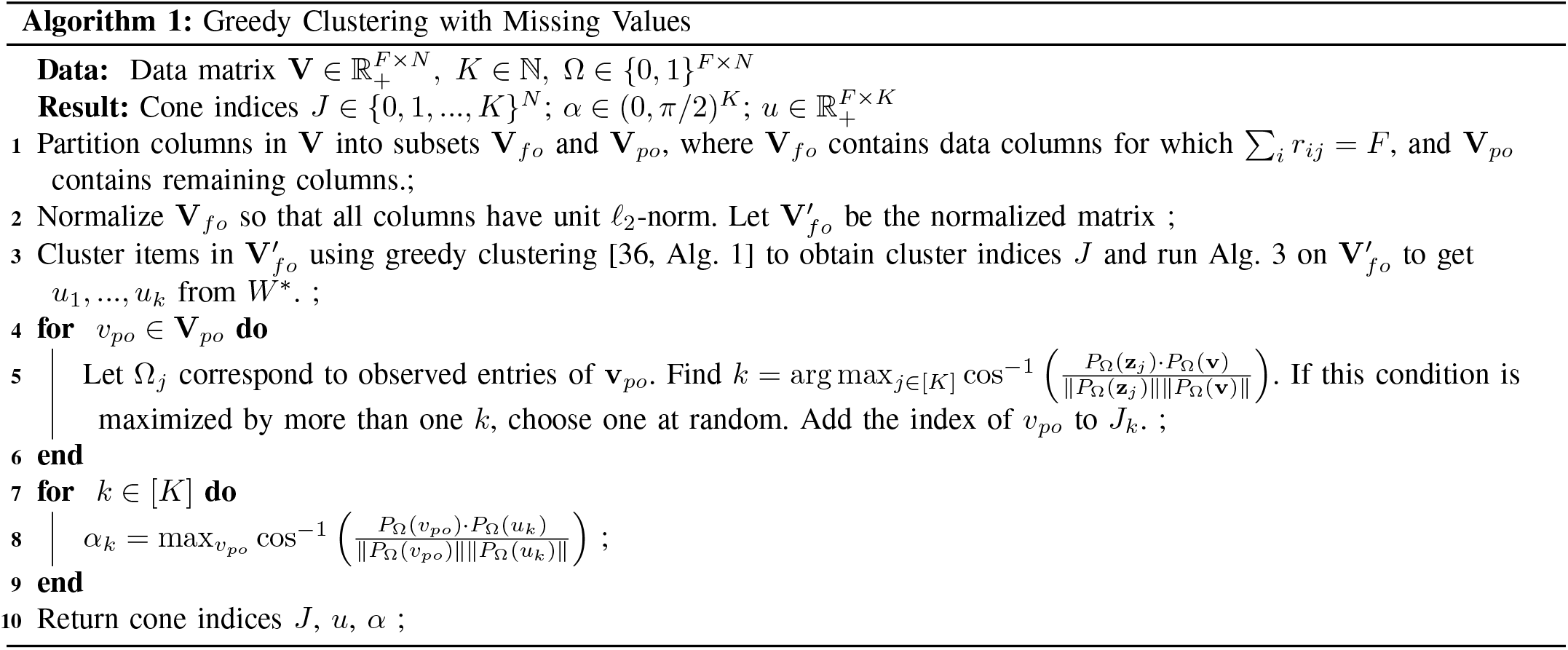

### Lemma 1

(Greedy clustering with missing values). *Let* Ω *indicate the missing values of v_po_. Let α_k_ be the defining angle of C_k_ and P*_Ω_(*α_k_*) *be the defining angle of the cone resulting from projecting C_k_ onto the missing value plane from* Ω. *If, for exactly one k*,

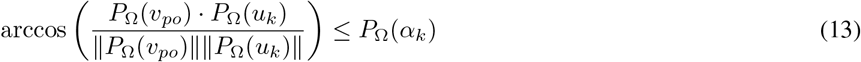

*then v_po_ originated from the corresponding C_k_. If α_k_ are identical for all k, Alg. 1 will cluster v_po_ correctly*.

*Proof*. The result follows directly.

Now consider feasibility of imputing data points using the 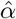 and 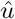 from Alg. 2. Clearly, the missing values plane for each point intersects the original corresponding cone defined by the true *u* and *α* of the cone. We know the 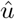 fall somewhere within the original cones, but if the 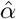 are too small, the new cones may not intersect with the missing values plane.

### Lemma 2

(Feasibility of imputation algorithm). *The estimator in* (5) *is able to find an imputation within the feasible set given α*_1_,…, *α*_K_ *and u*_1_,…, *u_k_ returned by Alg. 1*.

*Proof*. Let vector *v_po_* be a partially observed version of *v_fo_* ∈ **V**. We define the angle between *v_po_* and cluster center *u_k_* in the *F*-dimensional space:

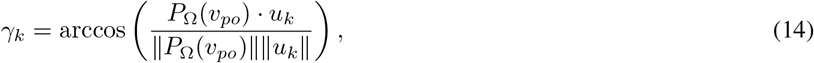

and between *v_po_* and the projected cluster center in the projected (*F – f*)-dimensional space:

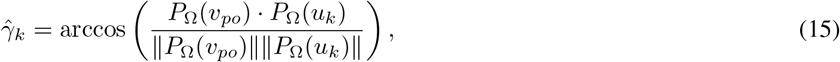

where Ω is the observed values indicator corresponding to *v_po_*. Then 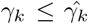 since *P*_Ω_(*v_po_*) · *u_k_* = *P*_Ω_(*v_po_*) · *P*_Ω_(*u_k_*) and ||*u_k_*|| ≥ ||*P*_Ω_(*u_k_*)||. Thus 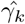 is large enough that an imputation on the missing values plane is feasible for each *v_po_*. Since *α_k_* = max *γ_k_*, all partially observed points labeled as belonging to *C_k_* can be imputed.

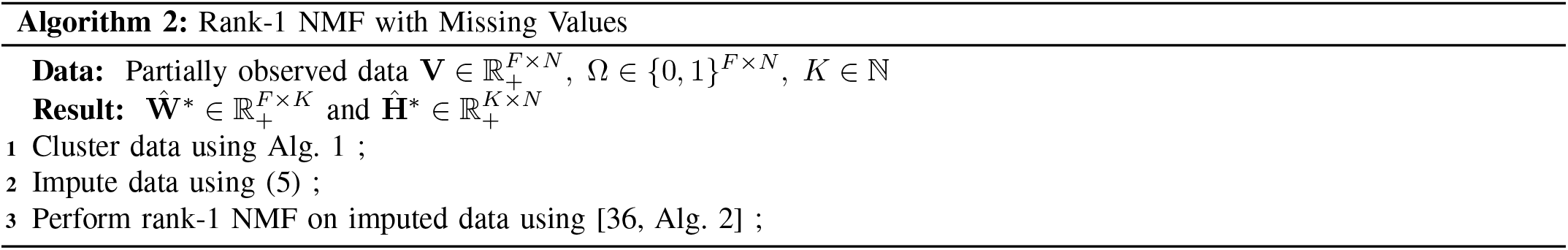

We extend bound (10) on the relative NMF error to missing values (Alg. 3). Note that the original bound allows for overlapping cones and does not assume (9) holds. It only requires all points be within *α_k_* of *u_k_*, which essentially allows the normalized perturbation matrix **E**_*k*_ to be upper-bounded by sin *α_k_*. If the missing entries of each *v_po_* are imputed using Alg. 1, then the perturbation from the original *u_k_*, which we denote 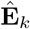, will be at most 2**E***_k_*. We can prove this using a worst-case scenario.

### Theorem 1

(Rank-1 NMF with missing values). *Suppose* **V** *is drawn from K cones and missing values are introduced to get **V***_po_*. If Alg. 2 correctly clusters data points and Alg. 1 is used to perform imputation, then*

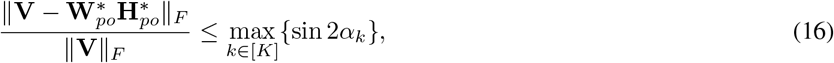

*where* 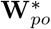 *and* 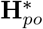 *are found by Alg. 3*.

*Proof*. Suppose there are two points *v*_1_ and *v*_2_ in a cone, as indicated by the solid circle in Fig. 4. Then *u* will be at an angle *α* from both *v*_1_ and *v*_2_. Now suppose *v*_2_ contains missing values. Then the new *v*_1_ will be the only vector in the cone, *v*_2_ is imputed using (11), where 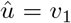, and 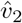 is at an angular distance sin 2*α* from 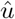. (One can check that if there are more than two points in the cone, this distance cannot increase.) A worst-case imputation places 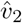 at an angle 2*α* away from *v*_1_ (suppose the optimizer places *v*_2_ at an angle greater than 2*α* from *v*_1_, but this is a contradiction since then *v*_2_ would be a better estimate than the optimum). The dashed circle in Fig. 4 represents points at an angle 2*α* from *v*_1_. Any 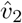 outside the dotted circle is at an angle greater than 2*α* from *v*_2_. So the shaded region indicates when the error may be greater than sin 2*α*. But the missing values of *v*_2_ allow for “movement” only along the axes. Since the intersection of a hyperplane with a cone is a finite-dimensional ellipsoid [41], [42], which is compact [43], *v*_2_ cannot “travel” via imputation to the shaded region without crossing a feasible region less than 2*α* from 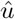. Hence the theorem holds and is tight.

**Fig. 4.**
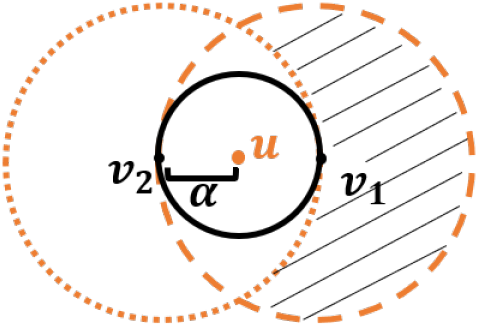
Geometric proof of relative NMF error bound.

## V. Probabilistic error

We now make some probabilistic assumptions on our data and missingness patterns to calculate the expected maximum error of optimal recovery imputation. First, consider a cone *C* in an *F*-dimensional space defined by *u* and *α*. Let us ignore the length of the vectors in *C* and preserve only the angles of the vectors from *u*. We can then represent vectors of an F-dimensional cone as points in an (*F* − 1)-dimensional ball. For example, a 3-dimensional cone can be represented as points in a circle, as in Fig. 4.

Let there be *N* points {*x*_1_,…,*x_N_*} ∈ ℝ^*F*^, drawn uniformly at random from *K F*-dimensional balls, labeled *B*_1_,…,*B_K_*. Let *d*(*x_i_, x_j_*) be the Euclidean distance between *x_i_* and *x_j_*. We assume there is at least one data point in each ball, and that the distance between any two points in a ball *B_k_* is less than the distance between any point in *B_k_* and a point not in *B_k_*. That is, for any *i, j* ∈ [*N*], *i* ≠ *j*,

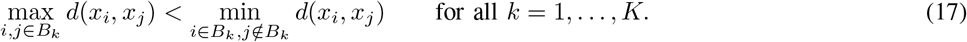

This is equivalent to the geometric assumption in (9), and we can correctly cluster any points drawn from such balls using Alg. 1. After obtaining the clusters, we can compute the minimum covering sphere (MCS) on the points in each cluster [44] (Fig. 5). This gives us *K* balls with *N_k_* points in each ball.

**Fig. 5.**
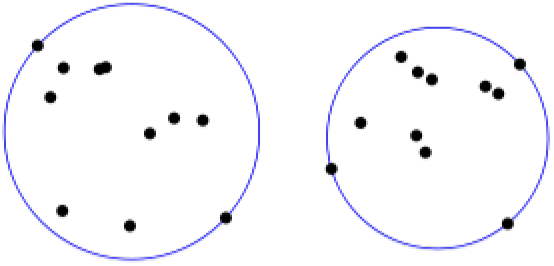
Minimum covering sphere in two dimensions.

Now suppose that we have partially observed entries in our data. Let the missingness of a point be a Bernoulli random variable with parameter *γ*. That is, *x* is fully observed with probability *γ* and partially observed with probability 1 − *γ*. There is now some uncertainty about the position of partially observed data points, so we will find the MCS for only the fully observedpoints. This is analogous to step 3 in Algorithm 2. By calculating the expected change in the radius of the MCS, we can calculate the expected change in its corresponding cone.

### Theorem 2

(Probabilistic bound on NMF error). *Given the setting described above, and assuming that the N points are drawn uniformly at random from the K balls, then after imputing with Alg. 1, we can tighten the bound in* (16) *to*

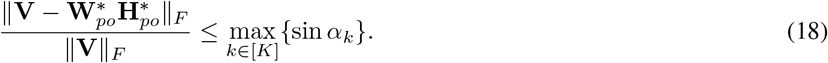

*Proof*. If the *N* points are drawn uniformly at random from the *K* balls, then 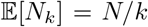, and the expected number of fully observed and partially observed points in each cluster is

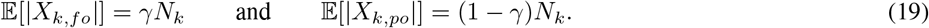

Clearly, the volume of the MCS can only decrease as |*X_k,fo_*| decreases. Let *R_max_* be the radius of MCS if there were no missing values, and let 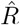 be the radius of the MCS of only the fully observed points. Then 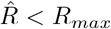 only if any *x* ∈ *X_po_* originally lay on the surface of MCS_*k,fo*_. Suppose the points are randomly distributed along the radius of the *F*-ball and we pick points to be partially observed uniformly at random. Let

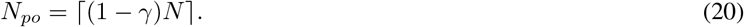

Assume *x_i_* are i.i.d. and uniformly distributed (without loss of generality) on [0,1]. This matches the assumption in the probabilistic analysis in [36] that the angles are drawn uniformly at random on [0, α] (see Fig. 6). Assuming a continuous distribution, almost surely no two points have exactly the same radius, and the probability of picking the *ℓ* outermost points is

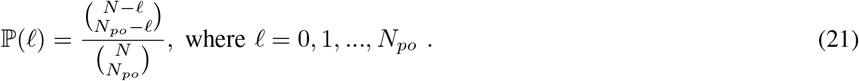

**Fig. 6.**
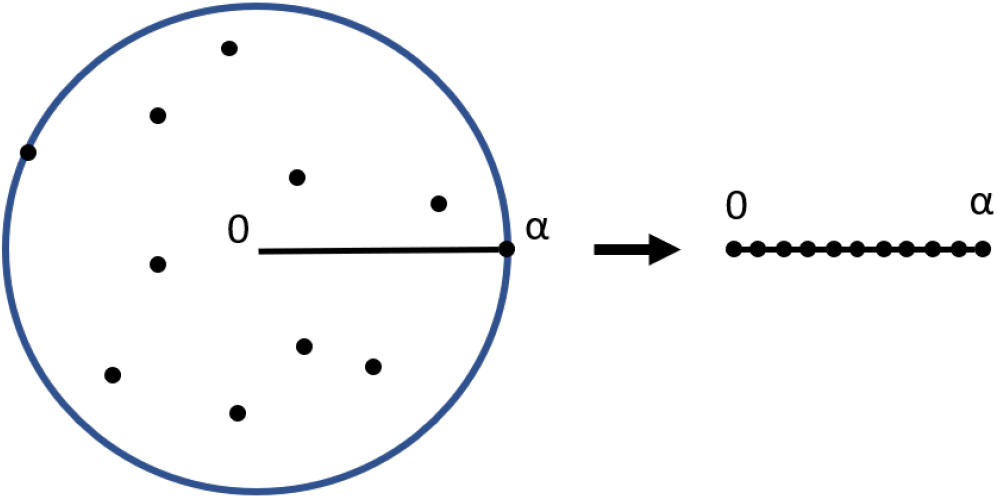
Assumption that points are uniformly random on the radius.

This gives us

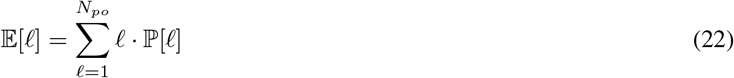

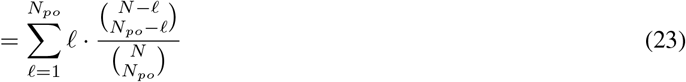

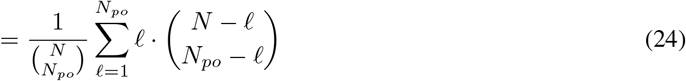

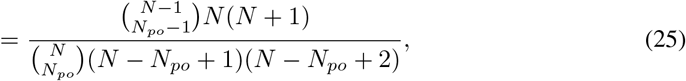

where *N_po_* is dependent on *γ*, as defined in (20).

The radius of the resulting MCS is dependent on the distribution of points along the radius. We can determine 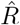 using order statistics. If we assume uniform distribution between 0 and 1, and order the points *x*_1_,…, *x_n_* so that *x*_1_ is closest to the center of the sphere and *x_n_* is farthest, the radius of the *n*th point, *R_n_*, is given by the beta distribution

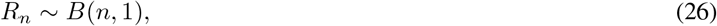

and

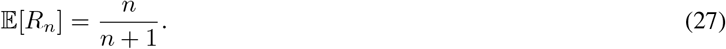

Thus if *ℓ* of the outermost points are chosen to be missing,

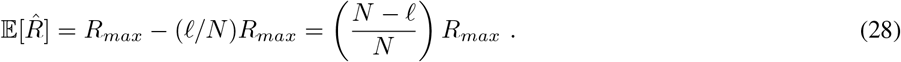

We illustrate with an example in Fig. 7. We can substitute 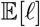 for *ℓ*, and since 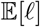 is a function of *γ*, we have derived the expected radius of the MCS as a function of missingness:

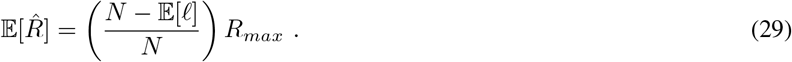

**Fig. 7.**
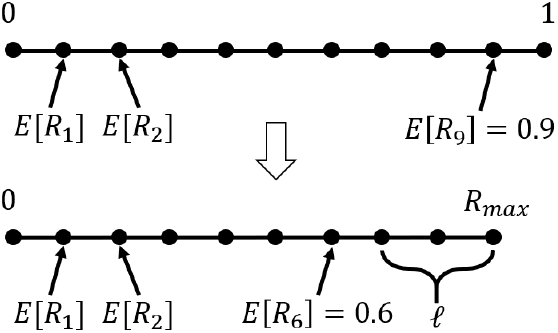
Example of 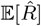 with *N* = 9 and *ℓ* = 3.

Now we reverse the arrow in Fig. 6. Due to the random distribution of points in the sphere, removing the *ℓ* outermost points does not change the expected center *u* of the MCS. Transitioning from spheres back to cones, we get

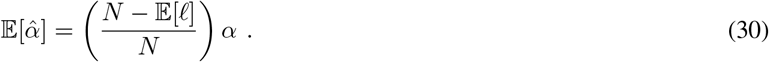

Thus

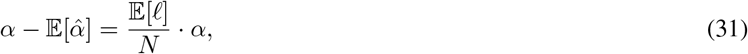

and the normalized Frobenius distance between 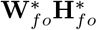 and **W**\***H**\* for a single cone is:

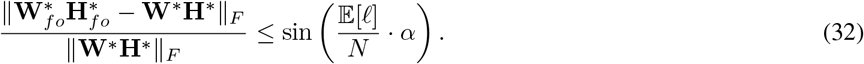

If we assume *v_n_* ∈ **V** are MCAR, the statistical mean of **V**_*fo*_ is the same as that of **V**. Since *v_n_* are uniformly distributed, the range of *v_n_* remains centered on the mean, so the expected center of the MCS does not change. Thus the maximum difference between a point *v* ∈ *C_k_* and its imputed point 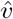 is sin *α_k_*, and the theorem follows.

### A. MCS with a different assumption

If instead we assume points are uniformly distributed in the volume of the ball, we find the change in radius as follows. First, calculate the volume of a *F*-dimensional ball of radius *R* = 1:

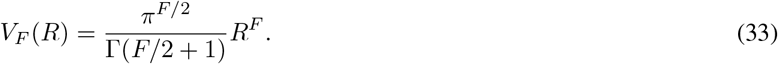

Then we calculate radius 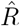 of an *F*-dimensional ball as:

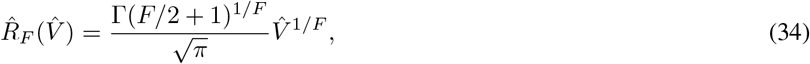

where volume 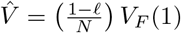.

The probability that a point *x* is in MCS_*po*_ is

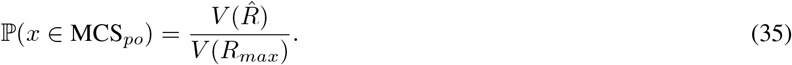

Thus the expected radius given a missing parameter *γ* is given by

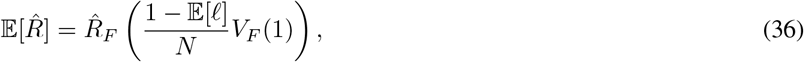

where 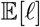 is a function of *γ*, and the expected NMF error is

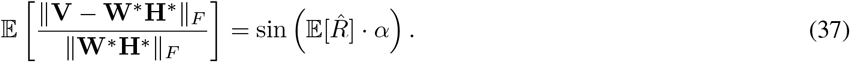

### B. Minimum covering spherical cap for normalized data

If the data is normalized such that each vector has an *L*_2_ norm of 1, all the points will fall on the surface of a sphere. Let there be *N* points {*x*_1_,…, *x_N_*} ∈ ℝ^*F*^, drawn at random from *K F*-dimensional spherical caps of a radius *R F*-ball, labeled *C*_1_,…, *C_K_*. Let *d*(*x_i_, x_j_*) be some distance between *x_i_* and *x_j_*. Assume there is at least one data point in each spherical cap, and that (9) holds.

The area of an *F*-dimensional spherical cap is

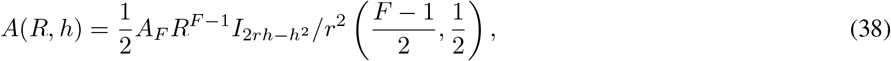

where 0 ≤ *h* ≤ *R, A_n_* = 2*π*^*n*/2^/Γ[*n*/2] is the area of the unit n-ball, *h* is the height of the cap, which can be calculated as a function of the angle *α* between the center and the edge of the cap, and *I_x_*(*a, b*) is the regularized incomplete beta function. Using the same style of analysis from the previous section, we can find the expected angle 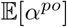 given a parameter *γ* for partially observed points. Thus,

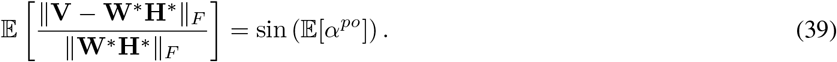

## VI. Experimental results

To test our algorithm, we first generate conical data satisfying the geometric assumption, using *N* = 10000, *F* = 160, and *K* = 40. We choose squared length of each *v* as a Poisson random variable with parameter 1, and we choose the angles of *v* uniformly. We then let **V** be partially-observed with Bernoulli parameter *ξ* to obtain **V**_*po*_. That is,

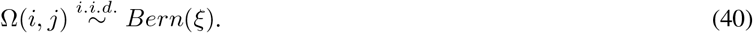

We run tests using *ξ* ∈ {0.4,0.55,0.7,0.8,0.9} and find imputation relative error for NMF:

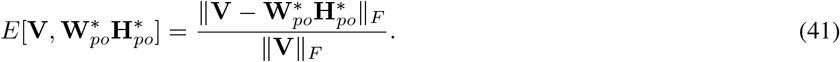

Fig. 8 shows relative error of our optimal recovery imputation with different values of *α* when we enforce correct clustering. The error for all *α* values and missingness percentages lies within the bound given by (16). Note that because our data is drawn uniformly at random, the error does not approach the worst-case bound.

**Fig. 8.**
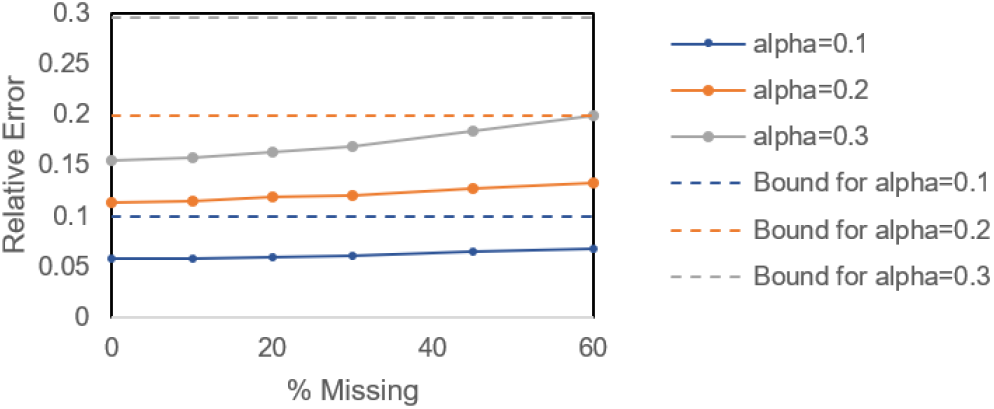
Relative NMF error of imputed conical data with correct clustering.

In the next experiment, we impute the conical data with *α* = 0.1 with other local imputation algorithms, including kNNimpute [45] with Euclidean, cosine, and Chebyshev (*L*_∞_) distances and iterated local least squares (itrLLS) [46]. We perform two tests with optimal recovery: one with enforced correct clusterings and one without prior knowledge of the correct clusterings. We use *α* = 0.1 and do not enforce correct clustering for Alg. 3 as before (see Fig. 9). We find *k* = 8 neighbors gives us the best results. Optimal recovery performs much better than other methods when clusters are known, and it performs similarly to other methods when they are not.

**Fig. 9.**
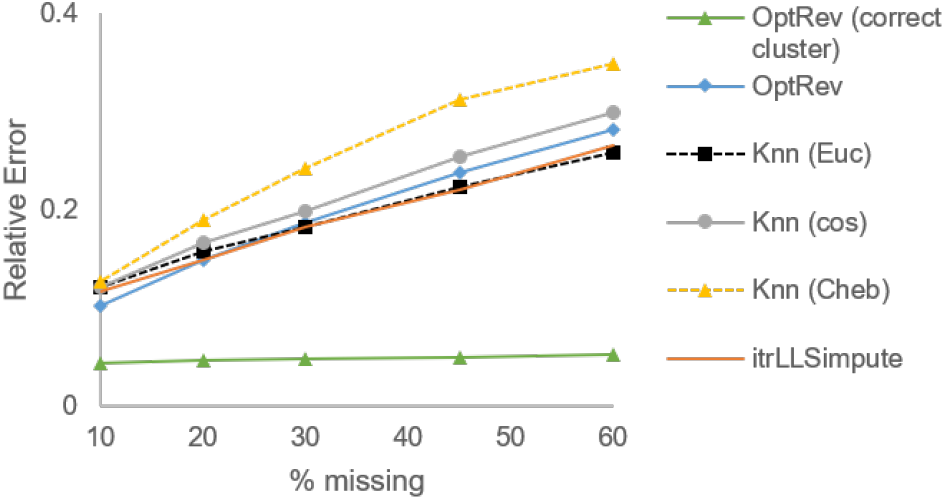
Relative NMF error for Conical data.

Following [36], the next experiment tests a subset of the hyperspectral imaging data set from Pavia [47]. We crop the 103 images to have 2000 pixels per image, set *K* = 9, corresponding to the different imagery categories, and introduce missing values in the same proportions as before (see Fig. 10). We also run tests with mice protein data [48] (see Fig. 11). The original dataset contains 1077 measurements with 77 proteins. We remove the 9 proteins that had missing measurements, then introduce missing values. We find *k* = 5 neighbors gives us the best results for kNNimpute on these datasets. On the mouse data, we also test bicluster BPCA [30] in addition to the other methods. The conical and Pavia test data were not sufficiently well-conditioned to run bicluster BPCA. See Tab. I for a comparison of run times. Our results demonstrate that optimal recovery performs similarly to kNN methods when clusters are not known beforehand. When clusters are known, optimal recovery performs similarly to more advanced methods (itrLLSimpute and biBPCA) in a fraction of the time.

**Fig. 10.**
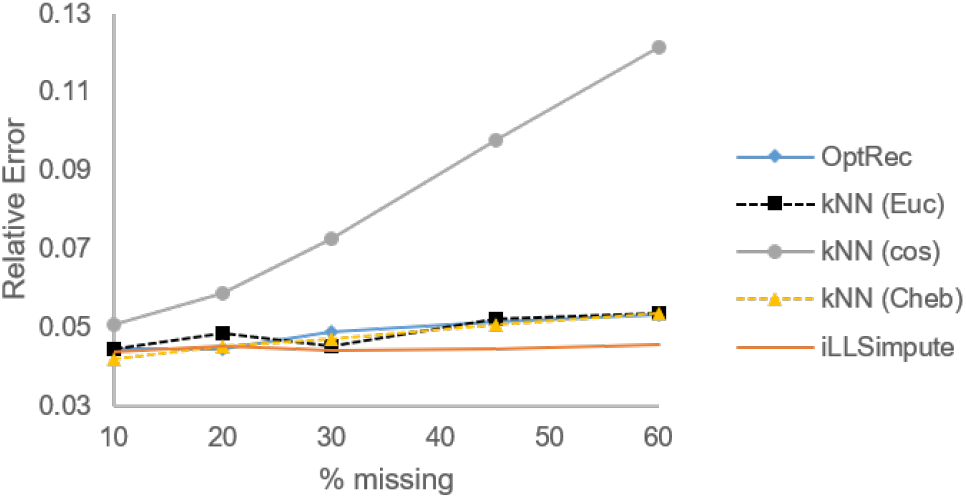
Relative NMF error for Pavia data.

**Fig. 11.**
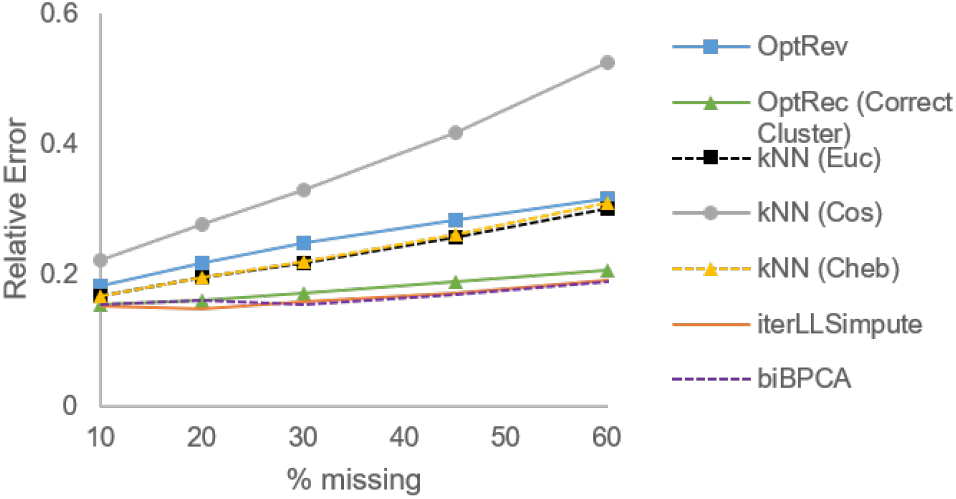
Relative NMF error for Mouse data.

**TABLE I.** Average imputation times for Mouse data in seconds.

| % Missing | 10 | 20 | 30 | 45 | 60 |
| --- | --- | --- | --- | --- | --- |
| **OptRec** | 0.51 | 0.48 | 0.63 | 0.91 | 1.41 |
| **OptRec (Correct Clusters)** | 0.48 | 0.48 | 0.58 | 0.85 | 1.36 |
| kNN (Euc) | 0.11 | 0.17 | 0.29 | 0.34 | 0.51 |
| kNN (Cos) | 0.10 | 0.15 | 0.19 | 0.27 | 0.41 |
| kNN (Cheb) | 0.08 | 0.16 | 0.23 | 0.35 | 0.52 |
| itrLLSimpute | 43.9 | 30.4 | 25.7 | 20.5 | 14.5 |
| biBPCA | 5000+ |  |  |  |  |

## VII. Conclusion

We have extended classical approximation-theoretic *optimal recovery* to the setting of imputing missing values, specifically for NMF. Fu et al. remark that “separability-based methods have many attractive features, such as identifiability, solvability, provable noise robustness, and the existence of lightweight greedy algorithms” [4]. We showed that imputation using optimal recovery minimizes relative NMF error under certain separability assumptions, and provided a straightforward algorithm for implementation.

Future work aims to extend optimal recovery to other settings of missing values in modern data science. Different applications (and therefore models) require different NMF algorithms and model assumptions. Thus different error bounds will be necessary [4]. On the experimental side, we plan to test our imputation algorithm on single-cell RNA sequencing data along with different clustering algorithms; these experiments will inform more specific heuristic refinements. We also aim to extend our algorithm to the scenario where complete cases for each cluster are not available.

